# The memory B cell response to influenza vaccination is impaired in older persons

**DOI:** 10.1101/2021.03.04.433942

**Authors:** Edward J Carr, Adam K Wheatley, Danika L Hill, Michelle A Linterman

## Abstract

Influenza imparts an age-related increase in mortality and morbidity. The most effective countermeasure is vaccination; however, vaccines offer modest protection in older adults. To investigate how ageing impacts the memory B cell response we tracked haemagglutinin specific B cells by indexed flow sorting and single cell RNA sequencing in twenty healthy adults administered the trivalent influenza vaccine. We found age-related skewing in the memory B cell compartment six weeks after vaccination, with younger adults developing haemagglutinin specific memory B cells with an *FCRL5*^+^ “atypical” phenotype, showing evidence of somatic hypermutation and positive selection, which happened to a lesser extent in older persons. We confirmed the germinal center ancestry of these *FCRL5*^+^ “atypical” memory B cells using scRNASeq from fine needle aspirates of influenza responding human lymph nodes and paired blood samples. Together, this study shows that the aged human germinal center reaction and memory B cell response following vaccination is defective.

**Summary:** Immune responses to vaccination wane with age. Using single cell RNA sequencing of influenza vaccine specific B cells, this study delineates changes in B cell memory generation, antibody mutation and their subsequent selection in older persons.

## Introduction

Influenza is a global health challenge. It has a large economic burden, estimated at $11.2 billion annually in the US alone (Putri et al., 2018). Around the world, there are an estimated 389 000 deaths a year attributable to influenza (Paget et al., 2019). Older individuals are particularly vulnerable to severe disease, with over two thirds of deaths among people aged 65 years and older (Paget et al., 2019), and are provided incomplete protection by vaccination, conferring an estimated 40% reduction in risk of influenza-like illness in the season following immunization (Demicheli and Rivetti, 2018). An understanding of what underpins the age-related decline in protection from the influenza vaccine is an important first step in developing rationally designed vaccines that are effective in the most vulnerable members of our communities (Pulendran and Davis, 2020).

The seasonal influenza vaccine is formulated annually to strains recommended by the World Health Organization Global Influenza Surveillance and Response System to combat antigenic drift and shift within circulating influenza strains (reviewed in (Hilleman, 2002)). Influenza infection is initiated when the influenza haemagglutinin (HA), a glycoprotein of the surface of the virion, binds sialic acid on host cells facilitating viral entry. A key correlate of protection after influenza vaccination is the production of anti-HA antibodies that block this interaction, and can neutralize the virus, stopping infection. The antibody response induced by seasonal influenza vaccines can be derived from multiple cellular pathways; from naïve and/or memory B cells being (re-)activated to form either early short-lived extrafollicular plasmablasts or to enter the germinal center (GC) (Tsang et al., 2014; Avey et al., 2020; Henry et al., 2019; Andrews et al., 2015; Turner et al., 2020). Within the GC, B cells undergo clonal expansion and somatic hypermutation (SHM) of the genes encoding the B cell receptor, followed by affinity-based selection and differentiation into long-lived antibody secreting cells or memory B cells that can provide protection against reinfection.

Healthy ageing is associated with a change in the composition and function of the human immune system, and impaired antibody production upon vaccination (Henry et al., 2019; Carr et al., 2016; Liston et al., 2016; Allen et al., 2020; Stebegg et al., 2020; Nakaya et al., 2015). In aged mice, the defects in antibody production after vaccination can be attributed to a poor germinal center response, as the early extrafollicular plasmablast response is comparable in adult and aged mice (Silva-Cayetano et al., 2020). However, whether there are age-dependent defects in the GC response upon vaccination of older persons has not been directly demonstrated. Here, we use single cell sequencing of HA-specific B cells after seasonal influenza vaccination to provide an in-depth understanding of the phenotype, clonality and hallmarks of GC selection of vaccine reactive B cells in older persons. We find the preferential expansion of a population of HA-binding atypical memory phenotype B cells in younger adults (22-36 yo), that does not occur to the same extent as in older people (67-86 yo). The atypical memory B cells show evidence of positive selection in the GC in younger, but not older, adults suggesting an impaired GC response in older persons. We confirm the GC origin of circulating atypical memory B cells upon influenza vaccination through clonal tracking of the progeny of human lymph node GC B cells in the blood. Together, these data provide evidence of an impaired GC reaction and memory B cell formation upon seasonal influenza vaccination in older individuals.

## Results and discussion

### Single cell RNA sequencing of haemagglutinin specific B cells, before and after vaccination captures the diversity of the influenza specific memory B cell population

To explore age-dependent changes in the humoral immune response a cohort of healthy volunteers received the 2016-17 trivalent seasonal influenza vaccine (TIV). Two age groups of volunteers were selected: 22-36 years old (n=10) or 67-86 years old (n=10). Venepuncture was performed immediately prior to vaccination (day 0) at day 7 and at day 42 after vaccination (Figure 1A), and sera and PBMC were collected to enable the analysis of antibody production and B cell subsets. All participants had virus neutralizing antibodies to the H1N1 A/California09 strain prior to vaccination, which were similar between age groups (Figure 1B). We observed a significant increase in day 7 titers in the 22-36 yo group, which was sustained at day 42 (Figure 1B). In the 67-86 yo group, there was a non-significant trend towards increasing titers of neutralizing antibody over the first week post vaccination, but this increased titer was short-lived. Similar reduced antibody titers after TIV have been shown in older individuals previously (Henry et al., 2019; Nakaya et al., 2015).

**Figure 1.**
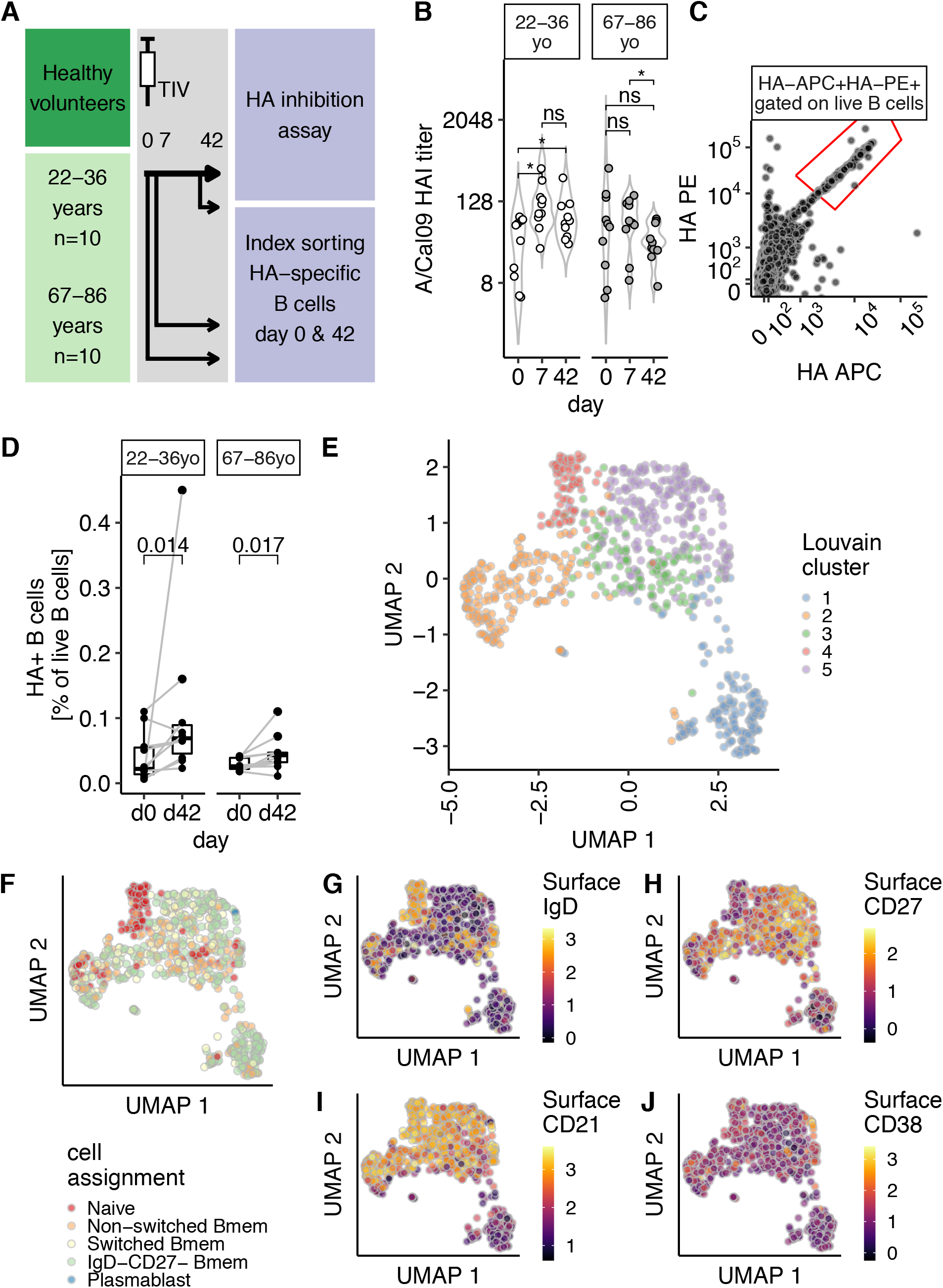
Single cell sequencing of haemagglutinin specific B cells to study the aged vaccine response. (A) Study design. Venepuncture performed on days 0 (just prior to immunization), 7 and 42. Peripheral blood mononuclear cells were isolated on the day of venepuncture and cryopreserved for later index sorting experiments. (B) Haemagglutinin inhibition (HAI) assay titers from days 0, 7 and 42 are shown for 22-36 year old or 67-86 year old volunteers as open or greyed circles respectively (n=10 in both groups). (C) Recombinant biotinylated haemagglutinin multimers conjugated with streptavidin-PE or - APC allows the identification of haemagglutinin specific B cells. Gated on live, singlet CD19+ lymphocytes. Full gating strategy is shown in Supplementary Figure 1A. (D) The proportion of haemagglutinin binding B cells increases after vaccination, for both 22-36 year olds and 67-86 year olds. Proportion expressed as percentage of live B cells, that did not bind free-streptavidin. (E) UMAP embedding of single cell RNA sequencing from A/Cal09-specific B cells, n=771 cells. UMAP projection based on the first 40 principal components using the features with the top 10% variance, after removal of low quality cells (Supplementary Figure 1) and size normalization by deconvolution. Louvain clustering reveals 5 clusters. (F) Cell identity assignment based on published transcriptional profiles of 29 human immune subsets including the following B cell subsets: IgD+CD27-‘naive’, IgD+CD27+ ‘non-switched Bmem’, IgD-CD27+ ‘switched Bmem’, IgD-CD27-‘IgD-CD27-’, IgD-CD27+CD38+ plasmablasts ‘PB’ and plasmacytoid DCs. The following numbers of cells were identified: naive 120 cells; non-switched Bmem 165 cells; Switched Bmem 144 cells; IgD-CD27-Bmem 334 cells; Plasmablast 3 cells and plasmacytoid DC 1 cell. No cells were defined as T cells or members of other lymphoid or myeloid lineages. (G-J) UMAP embedding as in (E), showing the logicle transformed index sort surface expression of IgD (G), CD27 (H), CD21 (I) and CD38 (J) proteins. In (B), *P* values from paired Mann-Whitney tests (on log2 transformed data) are summarized: *P*>0.05 ns; 0.05>*P*>0.01 *; 0.01>*P*<0.001 **. In (D), samples from the same individual are indicated with a grey line and *P* values shown are from a paired two-tailed Mann-Whitney test. In (G)-(J), the scales reflect the decimal log of the logicle transformed fluorescence intensity value.

To investigate how the composition of the memory B cell compartment is affected by age we index flow-sorted HA-specific B cells prior to, and 42 days, after seasonal influenza vaccination from all members of our cohort (Figure 1C, Supplementary Figure 1A). Whilst both age groups increased the number of HA-specific B cells after vaccination (Figure 1D, Supplementary Figures 1B-D), the fold change after vaccination was larger in younger (median FC 3.07) than in older adults (median FC 2.04). We sequenced 952 individual B cells using plate-based SMART-Seq (Picelli et al., 2014). After initial quality control (Supplementary Figures 2A-D), we retained 789 individual B cells. UMAP embedding revealed a small outlier population of 18 cells that had a gene expression profile consistent with plasma cells, with high expression of *PRDM1, IRF4* and *XBP1* (Supplementary Figures 2E-H). Analysis of their index sorting cell surface phenotype showed high levels of CD38 and low CD20 (Supplementary Figures 2I & J), with dim HA staining consistent with a plasma cell phenotype. These cells were excluded from downstream analyses to enable a focus on memory B cells. After these quality control steps, 771 B cells were used for analysis; and no single person dominated the dataset at either time point (Supplementary Figures 2K & L).

Within the transcriptome profiles of HA specific memory B cells, we found 5 clusters (Figure 1E), which could be separated from each other by several hundred putative marker genes (Supplementary Figures 3A & B). To determine whether these five populations correspond to known B cell subsets, we used published single cell transcriptional profiles of 29 different human immune subsets (Aran et al., 2019; Monaco et al., 2019) (Figure 1F). A single cell was called a plasmacytoid dendritic cell, whereas all others were classified as B cell subsets. To confirm this classification, we used our index sort information to confirm surface expression of selected proteins. Together, these suggested that UMAP cluster 4 (red, Figure 1E), were naïve B cells, being both surface IgD^+^ (Figure 1G) and surface CD27^-^ (Figure 1H). UMAP cluster 1 (blue, Figure 1E) was predominantly IgD^-^CD27^-^ Bmem by the transcriptional classifier (Figure 1F), and had low surface expression of CD21 protein (Figure 1I). The small number (3 individual cells) of transcriptionally assigned plasmablasts (Figure 1F) expressed high levels of surface CD38 (Figure 1J), suggesting a cellular snapshot of partial differentiation towards plasma cells. These single cell transcriptomes of HA-specific B cells therefore comprise both naïve B cells and further differentiated sub-populations, which are incompletely described by the classical IgD/CD27 gating of B cell populations.

To further assess the cellular identity of these UMAP clusters, we next examined putative marker genes for each cluster (Figures 2A-B, Supplementary Figures 3A and B). To compare gene expression between UMAP clusters we selected the 15 most significant comparisons between each cluster and any other (Figure 2A). We next assessed the three most discriminatory genes between each UMAP cluster and all others, alongside a supervised analysis with several biologically relevant B cell genes – activation markers (*FCRL5, CD86, TNFRSF13B* encoding TACI, *FCER2, CD27, CD24*) transcription factors (*TBX21* encoding Tbet, *BCL6*, *PRDM1*, *PAX5*), chemokine receptors (*CCR7*, *SELL*, *CXCR3*, *CXCR4*) and components of the DNA repair and SHM machinery (*XRCC6, AICDA*) (Figure 2B & Supplementary Figure 3C). UMAP cluster 1 (blue, Figure 1E) is characterized by the expression of *FCRL5, FCRL3, ITGAX* and *CD86* (Figures 2A-B & Supplementary Figure 3), and lack of *CCR7* and *SELL* (Figures 2A-B & Supplementary Figure 3). Whilst *TBX21* appears restricted to the *FCRL5*^+^ population (Figure 2B), pairwise comparisons for *TBX21* do not reach statistical significance (FDR adjusted *P*=0.15). This *FCRL5*^+^ population is analogous to a population of atypical memory B cells seen after malaria challenge (Kim et al., 2019), increased autoimmune disease (Rubtsov et al., 2011), increased in ageing (Rubtsov et al., 2011; Hao et al., 2011), and similar to the influenza vaccine responding cells described in younger persons (Horns et al., 2020). Naïve B cells, UMAP cluster 4, expressed high levels of characteristic markers of naïve B cells in other gene expression datasets, *TCL1A* (Nagumo et al., 2009), *YBX3* (Longo et al., 2009; Horns et al., 2020) and *IL4R* (Figures 2A-B & Supplementary Figure 2). Clusters 2, 3 and 5 are less distinct on UMAP embedding (Figure 1E), and are less clearly defined by single putative marker genes (Figures 2A and 2B). For clarity within the text and figures (Figure 2B), we refer to these clusters as *MALAT1*^+^ (FDR adjusted *P*=3.53×10^-16^), *LY9*^+^ (FDR adjusted *P*=1) and *MARCKS*^+^ (FDR adjusted *P*=0.23), respectively.

**Figure 2.**
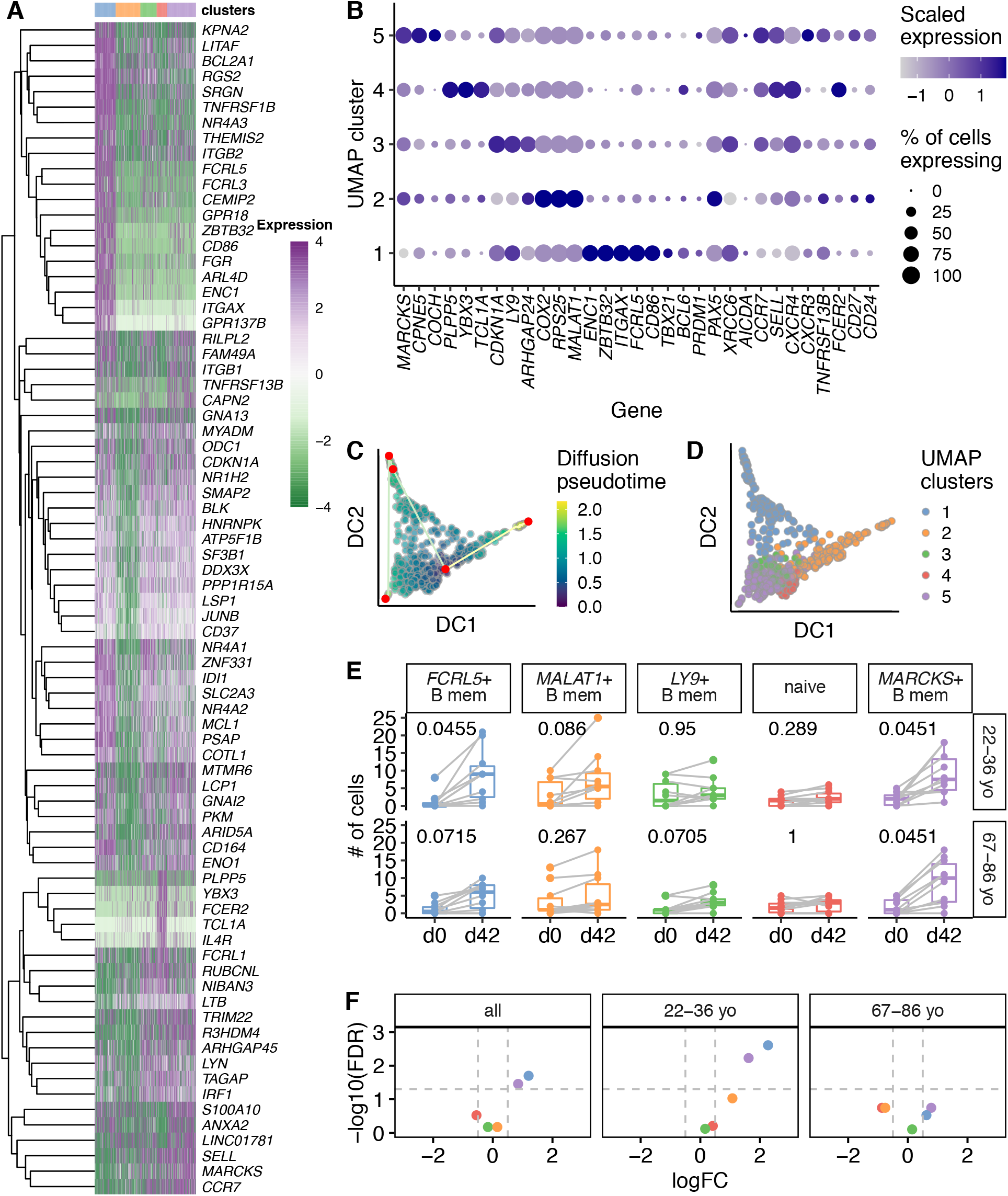
Transcriptional landscape is altered between young and old HA-specific memory B cells. (A) Heatmap showing expression of the top 15 features from t-tests distinguishing each UMAP cluster from *any* other cluster (FDR<0.01, log2 fold change>2). Each row is a feature (n=76) and its gene symbol is shown. Each column is a single cell (n=771). Cells are ordered by UMAP cluster, as shown in the color bar above the heatmap. The 15 features with the largest fold-changes were selected with tied positions allowed, and a feature could appear in more than one comparison. Features that did not map to a gene symbol and duplicated features were removed prior to plotting. Log2 expression values are row-normalized and centered. (B) Dotplot showing the expression of selected genes in each UMAP cluster. The size of the dot reflects the proportion of cells within that cluster which express the gene of interest. The color of each dot is scaled according to normalized expression of the given gene in that cluster. Genes were selected as follows: the top 2 genes from a t-test comparison between each cluster from *all* other clusters (L2FC>0.5, FDR<0.25); biologically relevant B cell genes - selected genes from (A), B cell transcription factors, DNA repair proteins, B cell chemokine receptors and other B cell surface receptors. (C) Diffusion co-efficient (DC) based pseudotime analysis from A/Cal09-specific B cells from day 42. Cells are shaded based on their position in pseudotime. Nodes are plotted in red and paths are shown by straight lines. (D) Pseudotime analysis as in (C), with colors determined by the UMAP clusters in (A) and defined in Figure 1. (E) Boxplots of the numbers of cells within each UMAP cluster comparing day 0 and day 42 cell numbers sorted for 22-36 year old and 67-86 year old individuals. Clusters are labelled with putative surface marker genes shown in (B). *P* values from 2 tailed paired Mann-Whitney tests are shown, after Benjamini-Hochberg correction for 5 tests. (F) Volcano plots from differential abundance analysis for the whole study (‘all’), or the two age groups individually. Shown are - log10 (Benjamini-Hochberg) FDR and log2 fold change (L2FC). Grey dashed lines are shown at - log10(0.05) and at L2FC±0.5.

To infer pathways of differentiation in circulating antigen-specific B cells we performed trajectory analysis (Figures 2C and 2D). Pseudotime starts adjacent to naïve B cells and progresses along two paths (Figure 2C). One path is towards the *MALAT1*^+^ cluster. The second path progresses through *FCRL5*^+^ cells and then to the three plasmablasts that are co-clustered with *MARCKS*^+^ cells, suggesting that *FCRL5*^+^ atypical Bmem cells differentiate from naïve B cells. This cellular differentiation is unlikely to be direct; it will occur after naïve cells home to the B cell follicle in the draining lymph node, and it will involve intermediate phenotypes – perhaps germinal center B cells – as we have sampled the circulating precursor and progeny populations in blood.

To determine whether the abundance of these different UMAP clusters was altered by vaccination and age, we examined the number of cells in each cluster from each individual at days 0 and day 42. Of all the HA cells sequenced, there was an increased number of *MARCKS*^+^ Bmem and *FCRL5*^+^ atypical Bmem cells after vaccination, although the increase in *FCRL5*^+^ atypical Bmem cells was not significant in older persons (Figure 2E). To adjust for the number of cells sorted at each time point, we performed differential abundance analysis (Figure 2F), confirming an increase in abundance of *MARCKS*^+^ and *FCRL5*^+^ cells in 22-36 yo and not in 67-86 yo after vaccination. These data indicates that the aged immune system fails to generate Bmem cells upon vaccination, including *FCRL5*^+^ atypical B cells that have previously been reported to accumulate in older individuals (Rubtsov et al., 2011; Hao et al., 2011).

### Haemagglutinin-specific B cell receptor repertoires display age associated differences and the expansion of IGHV1-69 clones with broadly neutralizing complementarity determining regions is favored by younger individuals

Having demonstrated that a dampened Bmem response occurs in aged individuals after immunization, we hypothesized that a corresponding alteration in the B cell receptors selected after TIV may also exist. Of 771 cells that were retained after transcriptome quality control, 662 gave Ig heavy chain sequences that were assigned V segments, of which 661 resulted in a productive, in-frame B cell receptor. Assessing the constant region of the heavy chain, we found the *MARCKS*^+^ and *FCRL5*^+^ populations were largely class switched to IgG or IgA, and naïve cells were predominantly IgM positive (Figures 3A & B). Whilst naïve cells express IgM and IgD transcripts, with surface IgD protein (Figure 1G), the IgM transcript has a higher expression per cell than IgD, favoring the assignment of naïve cells as IgM^+^ during *in silico* BCR assembly (Figures 3A & B). The distribution of V segment family usage at day 42 was altered between the two age groups (Figure 3C, *P*=0.002, by Fisher’s test), with *IGHV4* more frequently used in B cell receptor heavy chains in younger individuals. V segment family usage was not altered between B cell subsets as defined in by UMAP clusters (Figure 3D). We next assessed heterogeneity in the usage of V segments at the allele level, where we found age-related differences (Figure 3E, *P=*0.019 by Fisher’s test). *IGHV4-39* was enriched in HA-specific cells from 22-36 yo participants and has been previously reported to be a common V segment used by several broadly neutralizing antibodies against pdm1H1N1 (Thomson et al., 2012). The most common *IGHV* allele in both age groups is *IGHV3-23*, a V segment associated with polyreactive HA antibodies (Guthmiller et al., 2020). The second most common allele in both age groups is *IGHV1-69*, the V segment most commonly used in broadly neutralizing antibodies (bnAbs) to influenza A strains (Lingwood et al., 2012). *IGHV1-69* bnAbs have been described both after viral illness (Wrammert et al., 2011) and seasonal influenza vaccination (Nakamura et al., 2013). Several *IGHV1-69* bnAbs have solved crystal structures in HA conjugates (reviewed in (Chen et al., 2019)) and bind a conserved epitope on the HA stem. Importantly, the binding does not require light chain interactions (Lingwood et al., 2012). The consensus properties are phenylalanine at position 54, within the CDR2; a preceding hydrophobic residue at position 53 and a tyrosine at positions 97, 98, 99, or elsewhere within CDR3 (Avnir et al., 2014). From these parameters we predicted the bnAb capabilities of the *IGHV1-69*^+^ B cells within our study (Figure 3F). Intriguingly, these rare bnAb B cells appeared to expand in 22-36 yo after TIV but not in 67-86 yo (Figure 3G), with a tendency for the 22-36 year olds to expand *IGHV1-69* bnAb-expressing B cells with *FCRL5*^+^ atypical Bmem transcriptional profiles (Figure 3H). Further, these data support the observation that serial TIV administration in aged individuals may not expand stem-directed bnAb (Andrews et al., 2015). Together these data suggest that older individuals have an altered vaccine responding *IGHV* repertoire, either from changes in the repertoire available to respond to the vaccine, or in the subsequent selection of vaccine activated B cells.

**Figure 3.**
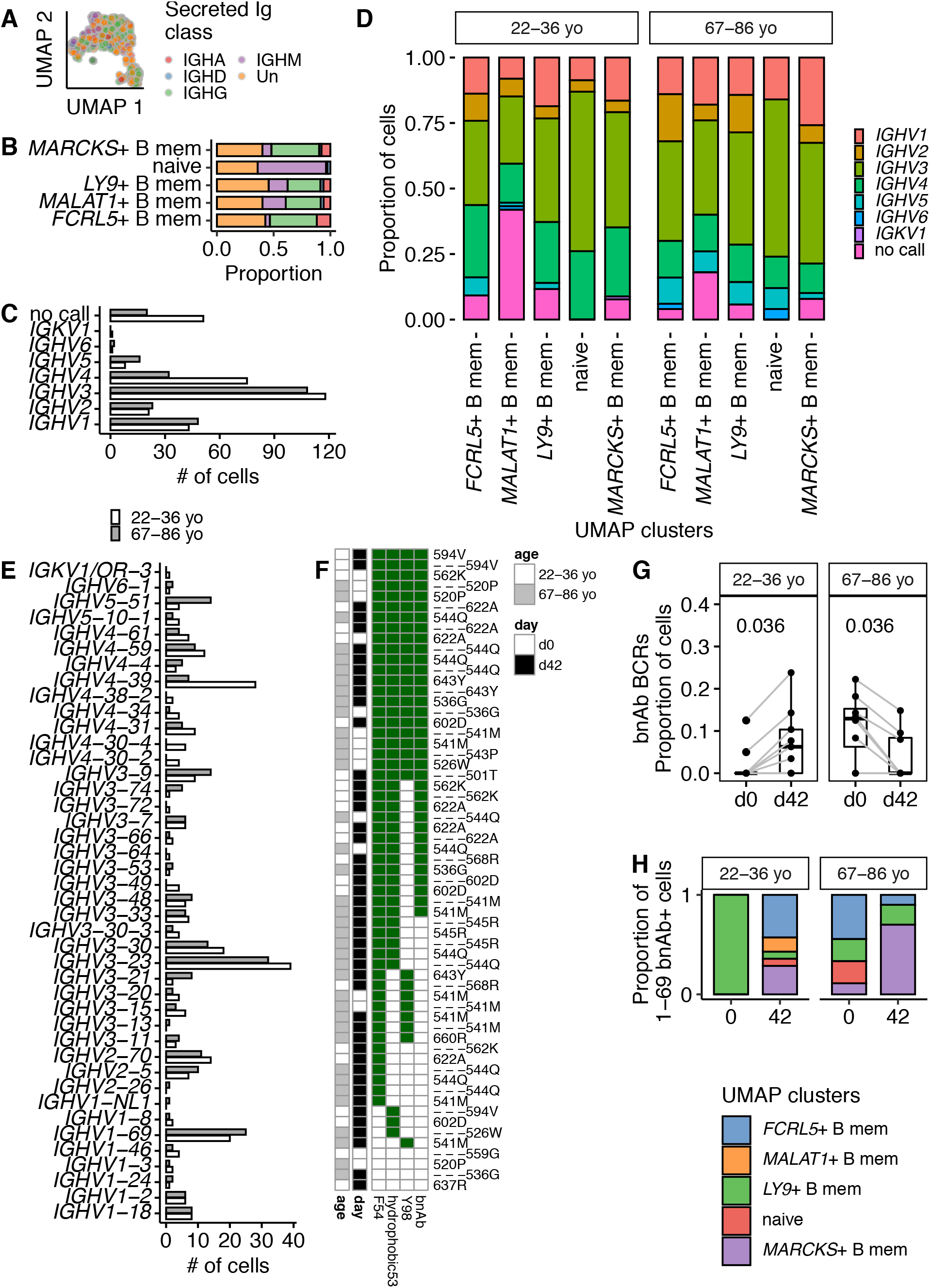
Differences in heavy chain V segment usage, including broadly neutralizing *IGHV1-69*, in aged individuals after TIV immunization in aged individuals after TIV immunization. (A) UMAP embedding as previously, showing secreted immunoglobulin classes of that cells’ most abundant immunoglobulin heavy chain transcript. Isotypes are combined to immunoglobulin class (e.g. IgG1, IgG2, IgG3, IgG4 grouped as IGHG): IGHA n=48 cells; IGHD n=12 cells; IGHG n=217 cells; IGHM n=112 cells; Un, unassigned n=272 cells. (B) For each UMAP cluster, the proportion of secreted immunoglobulin classes are plotted. Each immunoglobulin class is indicated with the same shading as in (A). (C) V segment family usage in the immunoglobulin heavy chain at day 42 for younger and older individuals. (D) V segment family usage by HA-specific immunoglobulin heavy chains at day 42 for younger and older individuals expressed as the proportion of the cells separated by each UMAP cluster. (E) V allele usage in the immunoglobulin heavy chain at day 42 for younger and older individuals. (F) Heatmap summarizing the presence of biophysical attributes characteristic of broadly neutralizing antibody for *IGHV1-69* B cells. The presence of F54, a hydrophobic residue at 53 and a tyrosine at 97, 98 or 99 are shown. An antibody is expected to be a bnAb if F54, there is a hydrophobic residue at position 53 and a Y within the CDR3. Presence of a characteristic is shown by a green box and empty boxes reflect its absence. On the left hand side of the panel, the age group of the cell is shown (cells from 18-36 year olds and 65-98 year olds in white and grey respectively) and the day of the sample is shown (day 0 and day 42 in white and black respectively). The individual identifiers are listed next to each row and some individuals. (G) The proportion of B cells from each study day that encode *IGHV1-69* bnAbs. Only those individuals with paired data from day 0 and day 42 are shown, which required > 1 successfully filtered B cell with a productive heavy chain at both days. BnAbs were defined as shown in (F). Grey lines link the same individual. Paired 2 tailed Mann-Whitney *P* values are shown. (H) The proportion of *IGHV1-69* B cells within each UMAP cluster for both age groups and on days 0 and 42 after TIV.

### The stringency of selection somatic hypermutation variants is reduced in older persons

A key site for diversification and selection of the B cell response is the germinal center which forms in secondary lymphoid tissues upon vaccination. Within the germinal center, B cells undergo SHM followed by affinity-based selection, enabling B cells with improved capacity to bind antigen to exit the germinal center as memory B cells or antibody secreting cells. The SHM frequency and selection of HA-binding B cells was assessed in all B cell subsets. In both groups there was an increase in the number of mutations in the heavy chain from day 0 to day 42 indicative of some of the circulating HA-binding B cells having participated in the germinal center response after vaccination (Figure 4A). At day 42, there were no differences at between 22-36 yo and 67-86 yo in the number of IgH mutations within framework (FR, Figure 4B) or complementarity determining regions (CDR, Figure 4C). These data suggest the processes of SHM are intact in vaccine responding B cells from in older recipients, similar to previous reports (Banerjee et al., 2002; Lazuardi et al., 2005; Dunn-Walters, 2016). To determine whether ageing influenced the positive selection of somatically mutated clones, we considered the replacement:silent (R/S) ratios in the heavy chain. The R/S ratio was higher in the CDR than the FR regions consistent with positive selection of mutations in the antigen-binding region (Mann-Whitney *P*=1.3×10^-35^, at day 42). There were no differences between age groups in the R/S ratios in the whole heavy chain (Figure 4D), or the FR (Figure 4E) or CDR regions (Figure 4F) at day 42 when all HA-specific B cell types were analyzed together.

**Figure 4.**
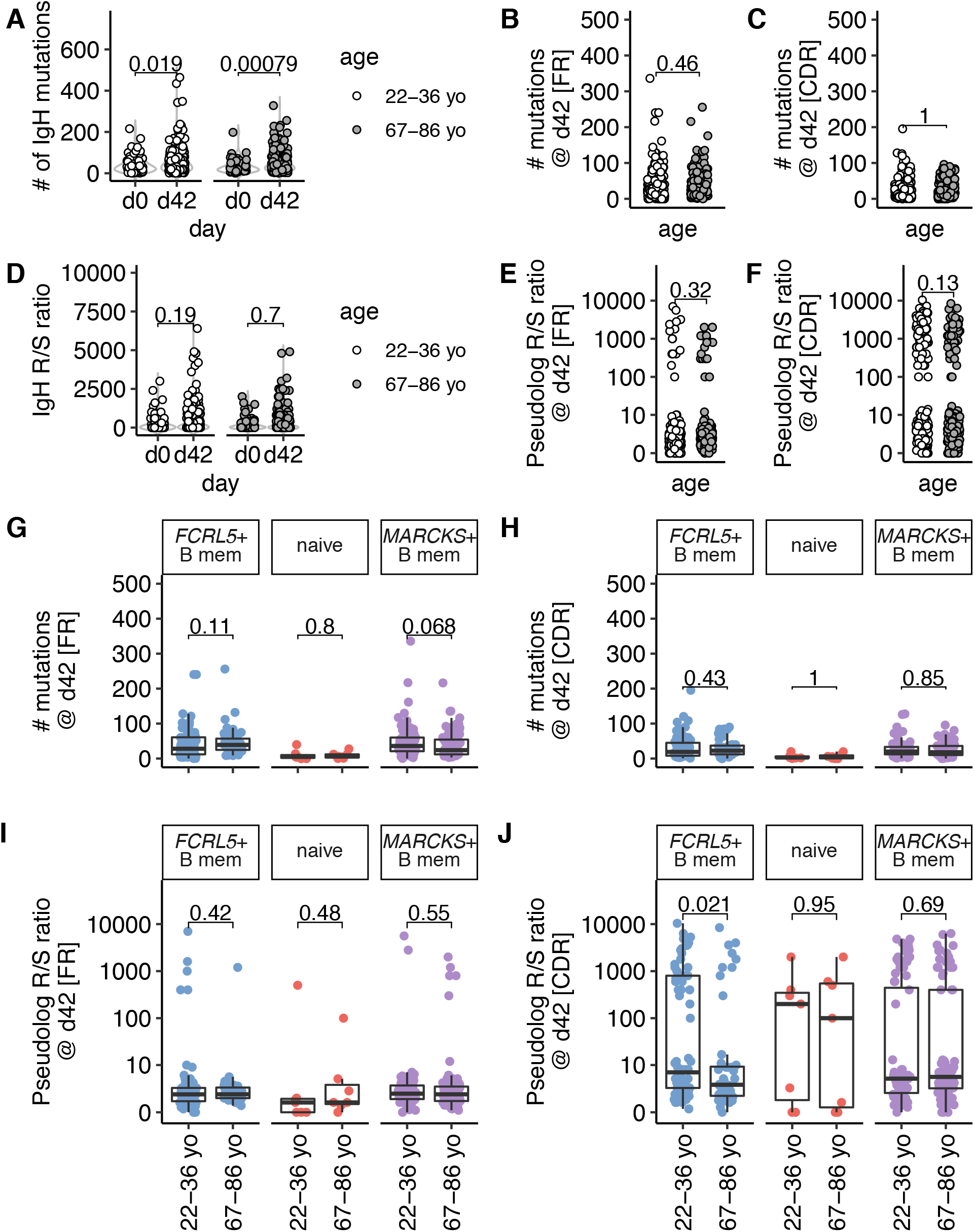
Somatic hypermutation is reduced in haemagglutinin specific memory B cells from aged individuals after TIV immunization. (A) The number of nucleotide mutations within the antibody heavy chain are shown for each cell for day 0 and 42 and for older and younger individuals. (B) and (C) The number of nucleotide mutations within the antibody heavy chain are shown for each region at day 42 and for older and younger individuals for FR (framework regions, B) or CDR (complementarity determining regions, C). (C) The ratio of replacement:silent mutations within the antibody heavy chain are shown for each cell for day 0 and 42 and for older and younger individuals. The replacement:silent ratio was calculated: # replacement mutations / (# silent mutations + 0.01), as many cells had zero silent mutations and >=1 replacement mutations. (E) and (F) The ratio of replacement:silent mutations within the heavy chain is shown for each cell from day 42 from both age groups. (G) and (H) The number of mutations in the antibody heavy chain at day 42 plotted by age group for each UMAP cluster, for FR (G) or CDR (H). (I) and (J) The ratio of replacement:silent mutations the antibody heavy chain at day 42 plotted by age group for each UMAP cluster, for FR (I) or CDR (J). In (G)-(J), each UMAP cluster is labelled and the colors correspond to its appearance in Figures 1–3. For (E, F, I & J), the ratio of replacement:silent mutations is calculated as in (D), and plotted as a pseudolog. In (A)-(J) P values from two tailed unpaired Mann-Whitney tests are shown. Where data are transformed for plotting (E, F, I & J), the test was performed on the untransformed data. The boxplots show the median, and inter-quartile range (IQR), with whiskers extending to the furthest data point, up to a maximum of 1.5x IQR. In (E, F, I & J), the boxplots correspond to the median and IQR of the transformed data.

To understand the mutation and selection of different HA-specific B cell subsets, we analyzed SHM and selection in the different UMAP clusters. We focused our SHM analyses on the two populations with the highest rate of class switch recombination, *MARCKS*^+^ and *FCRL5*^+^, with the naïve population as a control (Figure 3B). The highest numbers of mutations were found in *MARCKS*^+^ and *FCRL5*^+^ *MARCKS*^+^ and *FCRL5*^+^ clusters, with the lowest numbers of mutations found in the cluster of naïve cells, further supporting the assignments of cluster 4 as naïve B cell and *MARCKS*^+^ and *FCRL5*^+^ clusters as differentiated Bmem populations. We found no difference between age groups in the number of IgH mutations in the framework or complementarity determining regions in either Bmem subset (Figures 4G & H). Whilst the R/S ratios within the framework regions were unchanged between age groups (Figure 4I), there was a decrease in the R/S within the CDR in *FCRL5*^+^ B cells from older donors compared to younger individuals (Figure 4J), consistent with positive selection restricted to CDRs. This indicated that positive selection of this B cell subset after vaccination was impaired in ageing, similar to defective selection of GC B cells seen in Peyer’s Patches from older persons (Banerjee et al., 2002).

Further, these data suggest a germinal center origin of human *FCRL5*^+^ CD21^-^ atypical memory B cells after vaccination.

### FCRL5^+^ B cells are antigen experienced, germinal center emigrants

We have now identified an age-related diminution in the expansion of *FCRL5*^+^ HA-specific B cells after vaccination, and that these *FCRL5*^+^ cells have high a R/S ratio of their CDR mutations implying stringent BCR selection in younger persons. These attributes of selection and expansion of antigen specific B cells are hallmarks of passage through the GC, prompting the hypothesis that *FCRL5*^+^ Bmem cells are a circulating output of the GC. To formally test whether *FCRL5*^+^ Bmem cells are GC progeny requires paired sampling of the draining lymph node and blood at various timepoints post vaccination, with single cell transcriptomics to both identify GC B cells and retrieve BCR sequences to fate-map GC B cells into their circulating daughter populations. We therefore re-analyzed the Turner *et al*. dataset (Turner et al., 2020), in which a healthy 29-year-old man underwent serial fine needle aspirates (FNA) of his axillary lymph node with paired venepuncture, immediately before and days 5, 12, 28 and 60 after receiving a quadrivalent influenza vaccine (QIV). The FNA, PBMC and PBMC enriched for Bmem cells (IgD-enriched PBMC) samples from each day underwent single cell sequencing including B cell VDJ sequencing. First, all FNA timepoints were analyzed and, within the B cell compartment (defined in Supplementary Figures 4A & B), GC B cells were identified (Figure 5A & Supplementary Figure 4C). GC B cells were ~3% of all B cells within the lymph node FNA (Figure 5A). We used the B cell receptors of *bona fide* day 12 GC B cells to find daughter B cells in the circulation and lymph node (Figure 5B). Around 3% of all day 12 lymph node B cells were found to share a BCR with a GC B cells (Figure 5B), recapitulating the transcriptome profile analysis (Figure 5A). Circulating GC progeny were only found in the IgD-memory enriched PBMC samples and not in un-enriched PBMC (Figure 5B). Next, we identified B cells within the IgD-PBMC samples (Supplementary Figures 4D & E), and performed dimensionality reduction analysis of the circulating IgD-enriched B cells (Figure 5C). We identified 4 clusters, in the circulation, which we annotated based on the expression of B cell markers genes for naïve B cells, resting memory B cells, plasma cells and *FCRL5*^+^ Bmem (Figure 5D). Restricting our analysis to the circulating GC daughter cells identified in Figure 5B, we found that the circulating cells clonally related to the GC sampled at day 12 post vaccination were predominantly of an *FCRL5*^+^ Bmem phenotype (Figure 5E), with over 60% of all GC derived circulating B cells having a Bmem phenotype at day 28 after QIV (Figure 5F). These circulating *FCRL5*^+^ Bmem cells have high levels of SHM, favoring replacement CDR mutations (Supplementary figures 4F & G). This re-analysis demonstrates that *FCRL5*^+^ Bmem cells are GC derived, and that they are the most abundant circulating GC experienced B cell 28 days after immunization. Taken with our data showing fewer *FCRL5*^+^ Bmem forming in older persons (Figure 2E), that have impaired positive selection (Figures 4I & J), this indicates that the GC reaction upon vaccination is impaired in older persons.

**Figure 5.**
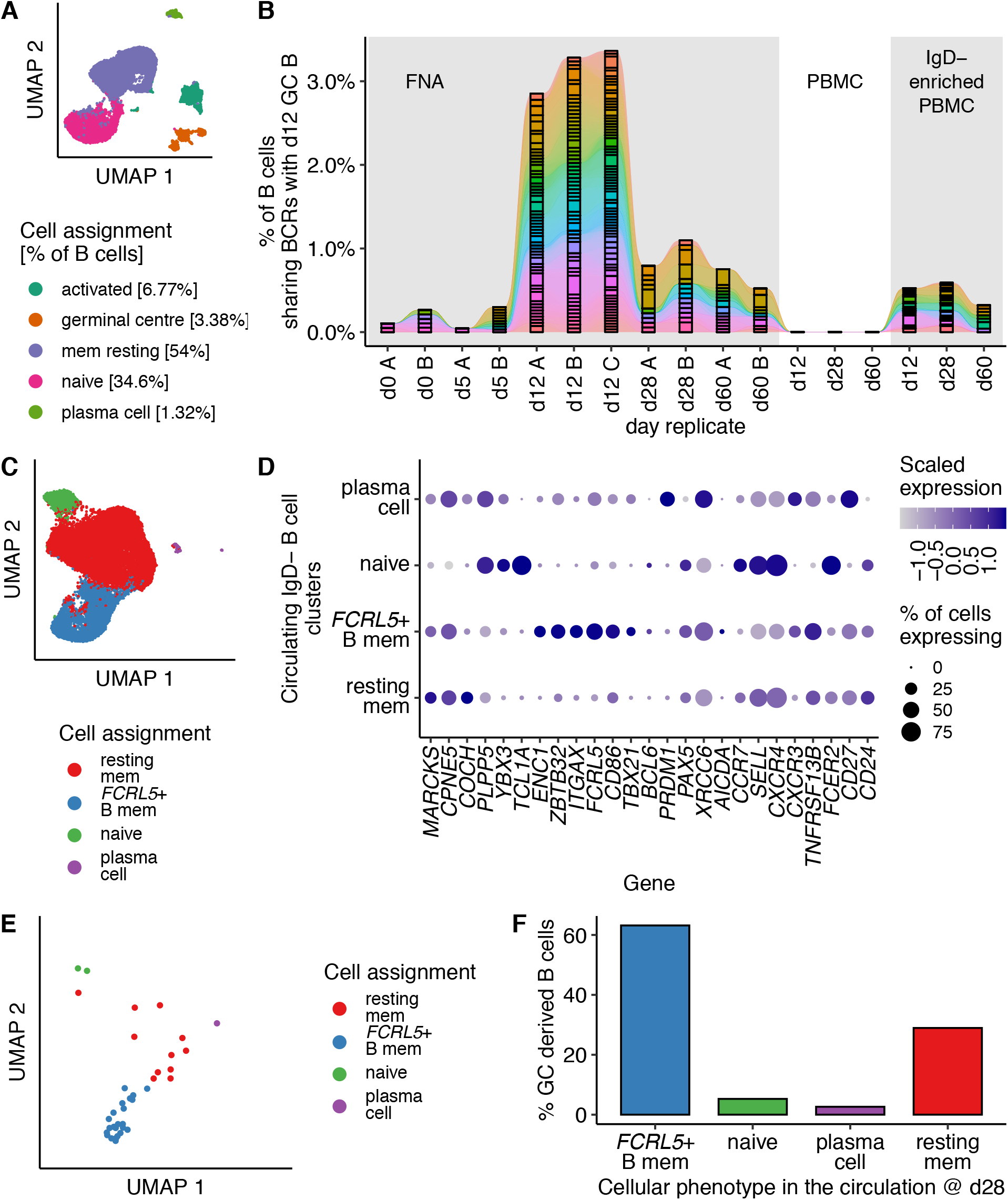
Germinal center emigrant memory B cells are *FCRL5*^+^. (A) UMAP of B cells (n=27,265 cells) from fine needle aspirates (FNAs) of draining axillary lymph nodes from a single healthy volunteer on days 0, 5, 12, 28 and 60 after quadrivalent influenza vaccine (QIV), as reported by Turner *et al*. (Turner et al., 2020). (B) The B cell receptors detected in germinal center (GC) B cells on day 12 after QIV immunization are shared with earlier LN B cells, and are detectable in peripheral blood mononuclear cells (PBMC) that have been enriched for B cell memory (IgD-) at days 28 and 60 post-vaccine. (C) UMAP of circulating B cells (n=21,568 cells) from IgD-enriched PBMCs at days 0, 5, 12, 28 and 60 after QIV. Clusters identified by Louvain clustering, and annotated based on (D). (D) Dotplot showing the scaled normalized expression of selected genes used to annotate the clusters identified in (C). (E) UMAP of circulating QIV-specific B cells from IgD-enriched PBMCs at day 28 which share a germinal center BCR, n=38 cells. (F) The percentage of QIV-specific B cells, present in the circulation at day 28, as in (E) is shown for each B cell cluster.

In this study we use the trivalent influenza vaccine and HA labeling to track vaccine specific B cells before and after immunization of younger people and elders. We show that the formation of Bmem cells is attenuated in individuals aged 67-86 yo, including the impaired formation of *FCRL5*^+^ B cells that have transcriptional signature characteristic of T-bet^+^ B cells (Li et al., 2016), previously termed ‘atypical memory B cells’. Similar populations are reportedly age-related memory B cells that are more prevalent in older donors (Rubtsov et al., 2011; Hao et al., 2011), exhausted B cells from chronic infections (Portugal et al., 2015; Moir et al., 2008), or from autoimmune diseases (Rubtsov et al., 2011). We used publicly available paired PBMC and FNA single cell sequencing to demonstrate *FCRL5*^+^ B cells are circulating progeny from the GC reaction. Recently, an *FCRL5*^+^ population has been described after malaria vaccination in naïve volunteers (Sutton et al., 2021), and our data supports this observation that *FCRL5*^+^ B memory cells can be part of a ‘typical’ immune response. In this study, atypical memory B cells from older persons showed evidence of SHM, but less evidence of positive selection, indicating impaired GC function in older people.

The process of positive selection in the GC requires the integration of GC B cell proliferation and mutation, followed by migration and selection. The GC is a polarized structure with two functionally distinct compartments - the light and dark zones (Allen et al., 2004; Bannard et al., 2013). In the dark zone GC B cells proliferate and the genes encoding the BCR undergo somatic hypermutation. Following mutation, dark zone GC B cells exit the cell cycle, and migrate towards the CXCL13-rich follicular dendritic cell (FDC) network of the light zone. In the light zone, GC B cells use their mutated BCR to bind antigen held on the surface of the FDCs in the form of immune complexes. A functional BCR enables the B cell to collect this antigen, process it and present it to a specialized subset of CD4+ T cells, Tfh cells, as peptide on MHCII. If the B cell can engage a cognate Tfh cell it will receive help, in the form of CD40L-dependent co-stimulation and cytokines. These Tfh-derived signals protect GC B cells from death and induce expression of cMyc, enabling the GC B cell to re-enter the cell cycle. GC B cells that do not receive appropriate help from Tfh cells die. The survivors migrate back to the dark zone and either undergo further rounds of proliferation and mutation, or differentiate into memory B cells or long-lived antibody-secreting cells that exit the GC (Mesin et al., 2016; Vinuesa et al., 2016). Thus, selection requires GC B cells to engage in productive interactions with both FDCs and Tfh cells. For this reason, age-dependent changes in B cells, FDCs and/or Tfh cells may underlie the altered selection observed in ageing.

There is evidence that the somatic hypermutation mechanism is intact in ageing, but that selection after mutation is impaired (Banerjee et al., 2002; Lazuardi et al., 2005; Dunn-Walters, 2016). This suggests that B cell extrinsic factors, such as FDC or Tfh cells, are responsible for the impaired selection in ageing. The formation of fully differentiated GC-Tfh cells is impaired in ageing, in both humans and mice (Lefebvre et al., 2012; Stebegg et al., 2020; Webb et al., 2021). Likewise, it has previously been reported that the activation of FDC upon antigenic challenge is impaired in ageing (Aydar et al., 2002) and that the FDC network in aged mice holds fewer immune complexes on its surface (Turner and Mabbott, 2017). Future work will be required to determine the relative contributions of changes in Tfh cells and FDC to the poor GC response in ageing.

## Materials and methods

### Trivalent influenza vaccination of healthy volunteers

Healthy volunteers aged 22-36 years old or 67-86 years old were immunized by the NIHR Cambridge BioResource. Individuals taking immunosuppressive drugs, or with active cancer were excluded, but we did not exclude individuals with controlled chronic conditions such as hypertension. Individuals underwent venepuncture (day 0) and then the seasonal trivalent influenza vaccine was administered intramuscularly (formulated in line with the World Health Organisation’s 2016-17 Northern hemisphere recommendations: i. A/California/7/2009 (H1N1) pdm09-like virus; ii. an A/Hong Kong/4801/2014 (H3N2)-like virus; iii a B/Brisbane/60/2008-like virus). Venepuncture was repeated on days 7 and 42. All venepuncture samples were 50mL and collected into silica-coated tubes (for serum, 4.5mL) or EDTA-coated tubes (for cells, 5 x 9mL tubes). All blood was collected in accordance with the latest revision of the Declaration of Helsinki and the Guidelines for Good Clinical Practice (ICH-GCP). Ethical approval for this study was provided by a UK local research ethics committee (REC reference 14/SC/1077). The Cambridge Bioresource has its own ethical approval (REC reference 04/Q0108/44). All volunteers provided written informed consent upon entry to the study.

### Peripheral blood mononuclear cell isolation and cryopreservation

Peripheral blood mononuclear cells (PBMCs) were isolated by density centrifugation over Histopaque 1077 (Sigma), counted and cryopreserved in 90% FCS / 10% DMSO (both Sigma). PBMC aliquots were frozen in a methanol bath (to freeze slowly ~ 1C/min) in a - 80°C freezer overnight. Once frozen, PBMC were transferred to liquid nitrogen for longer term storage. For the studies herein we used aliquots of ~10^7^ PBMCs.

### Haemagglutinin inhibition assays

Antibody titers pre and post vaccination were determined using the hemagglutination inhibition (HAI) assay using the standard WHO protocol, as previously described (Chen et al., 2010). Briefly, serum was treated with receptor destroying enzyme (RDE; Denka Seiken Co.) by adding 1-part serum to 3 parts RDE and incubating at 37 °C overnight, followed by RDE inactivation at 56 °C for one hour. Treated serum was then serially diluted 1 in 2 with PBS in 96 well v-bottom plates (Nunc) in 50 μL volumes. 4 HA units of A/California/04/2009 (H1N1) virus produced in vitro in MDCK cells was added per well, and incubated with serum samples for 30 minutes at room temperature. Then 50μL of 0.5% chicken RBCs (TCS Biosciences) in PBS was added per well, plates agitated manually, and incubated for 30 minutes. HAI titers were read manually as the reciprocal of the final dilution for which complete inhibition of agglutination was observed.

### Haemagglutinin specific bait for vaccine specific B cell sorts

Haemagglutinin from the A/California09 strain was previously modified with a single point mutation Y98F to minimize its binding to sialic acid (Whittle et al., 2014) (reducing non-specific binding to cell surface glycoproteins) and then biotinylated as previously (Wheatley et al., 2016). Biotinylated mutant Cal09-HA (1.5μg) was complexed with streptavidin conjugated to APC (8μL; Thermofisher #S868) or PE (8μL; Thermofisher #S866), in the presence of a protease inhibitor (1.5μL of the manufacturer’s recommended 100x re-suspension concentration; Merck #US1539131-1VL) in 115μL of phosphate buffered saline (PBS). The Cal09-HA, PBS and protease inhibitor were mixed first and then the streptavidin conjugates added in 5 steps (1.6μL each) every 15-20 minutes. The conjugated bait was stored in the dark at 4°C and used within 2 weeks, at 5μL / 10^6^ cells. We double-stained with streptavidin conjugates in two separate fluorophores for plate-based index sorting single B cells from the double-positive HA gate (Figure 1B; full gating strategy in Supplementary Figure 1A). A ‘free’ streptavidin conjugated to irrelevant fluorophore was used to gate out B cells that bound the streptavidin component of the bait.

### Index sorting of single cells with FACS after magnetic negative selection of B cells

Cryopreserved PBMC were collected from liquid nitrogen stores and thawed on wet ice. As soon as the last ice crystals thawed, 1mL of complete RPMI-1640, prewarmed to 37°C, was added to the cryovial and the whole contents transferred to 49mL of complete RPMI-1640 in a 50mL tube. For complete RPMI-1640, RPMI was supplemented with +10% heat inactivated fetal calf serum, 100U/mL penicillin and 100μg/mL streptomycin and L-glutamine. The cells were centrifuged and resuspended in complete media, pre-warmed to 37°C and allowed to rest for 1 hour at 37°C prior to viability counting. Untouched B cells were enriched using negative magnetic separation (eBioscience 8804-6867), following the manufacturer’s instructions with 2% FCS / PBS. Viable PBMC were counted with Trypan Blue (Sigma) exclusion and a hemocytometer.

Negatively selected B cells were filtered using a 30μm filter, counted and surface staining performed for 60 minutes in the dark at 4°C. Antibodies used are shown in table 1. Staining was performed in BD brilliant buffer (BD #563794), in the presence of human IgG as FcR block. Cells were washed in PBS and resuspended in 200-250μL of PBS for sorting.

**Table 1:** Antibodies used for single cell index FACS sorting.

| Marker | Clone | Fluorophore | Supplier | catalog number |
| --- | --- | --- | --- | --- |
| Viability dye | - | eFluor780 | eBioscience | 65-0865-14 |
| CD3 | UCHT1 | APC-eF780 | eBioscience | 47-0038-42 |
| CD14 | 61D3 | APC-eF780 | eBioscience | 47-9149-42 |
| CD16 | eBioCB16 | APC-eF780 | eBioscience | 47-0168-42 |
| HA-streptavidin bait | - | APC | In-house | (see methods) |
| HA-streptavidin bait | - | PE | In-house | (see methods) |
| CD21 | Bu32 | PE-Cy7 | BioLegend | 354911 |
| CD38 | HIT2 | BV421 | BioLegend | 303526 |
| CD20 | 2H7 | BV605 | BioLegend | 302312 |
| CD27 | O323 | BV711 | BioLegend | 302834 |
| CD71 | CY1g4 | FITC | BioLegend | 334103 |
| CD19 | SJ25C1 | BUV496 | BD | 564655 |
| IgD | IA6-2 | BUV737 | BD | 564687 |
| Free streptavidin | - | BUV395 | BD | 564176 |

Cells were index sorted on a BD FACSAria Fusion in cat 2 containment, with a 100μm nozzle into a 96-well plate (BioRad #HSS9641), containing chilled (4°C) RNA lysis buffer (SMART-Seq HT lysis buffer, Takara Clontech, #634439). This contained the oligo-dT primer. Immediately after the sort was completed, a labelled foil lid (Axygen, PCR-AS-200) was applied to the plate, it was spun briefly in a tabletop centrifuge and snap frozen in dry ice. Frozen plates were transferred to - 80°C for storage.

For each cell sorting batch, paired day 0 and 42 samples were processed from an individual from the 22-36 yo age group and an individual from the 67-86 yo age group. All 4 biological conditions were represented in each sorting batch. There were 10 sorting batches. Day 0 samples were sorted for 16 individual HA specific B cells, day 42 samples for 32 individual HA specific B cells (giving 16 + 16 + 32 + 32 = 96 cells / cell sort batch). Each biological condition was sorted in its own pre-prepared 96 well plate containing lysis buffer, and snap frozen at the end of that sort (rather than wait until an entire 96 well plate was filled), to minimize time spent at 4°C. Any extra HA specific B cells were collected on a fifth 96 well plate with any remaining lysis buffer.

Index sorting information was extracted from the FCS files using custom scripts based on the Bioconductor package *flowCore* (Ellis et al., 2019). Each sorted cell is given its own FCS file containing a single event. Our scripts extracted compensated, logicle transformed cell surface index data on a per cell basis, mapped to that cell’s eventual cDNA library (see next section). To analyze the flow cytometry phenotype of HA+ B cells in comparison to HA-B cells, the index FCS files, each containing a single event, were concatenated with their ‘master’ file into a combined file of all events from each biological sample. Flow plots are rendered with the *ggcyto* package (Van et al., 2018). Analysis of FCS files were performed using FlowJo v10 (Treestar).

### Single cell RNA sequencing using SMART-Seq

Samples were processed to cDNA in batches. Each batch comprised of 4 partially filled 96 well plates of flow sorted cells, from the same sort batch: paired d0 and d42 samples from one individual from either age group. cDNA was generated as per the manufacturer’s instructions (SMART-Seq HT kit user manual, version/date 121218, kit # 634436), using 20 PCR cycles. cDNA was synthesized using four 96 well BioRad T1000 PCR machines in parallel, with a single master mix used across all 4 plates. After cDNA synthesis, the 4 plates were consolidated into a single 96 well plate for ongoing processing (1 cell / well; the cDNA is not yet indexed; with a record of final well positions of each cell). If a cDNA synthesis batch was << 96 cells (this was predictable from the sort results), then additional cells from a fifth flow sorted plate were also processed and consolidated into the final 96 well plate. In this case, a fifth BioRad T1000 was used at the cDNA synthesis step.

In total, 10 x 96 well cDNA plates were generated. The 10 cDNA plates were cleaned-up using AMPure XP beads (Beckman Coulter #A63881), deep 96 well plates (Abgene #AB-0765) and a magnetic plate stand (ThermoFisher #AM10027) as described in the SMART-Seq HT manual (dated 121218). Cleaned cDNA was stored in 96 well plates (BioRad #HSS9641) at - 20°C. Quantification was performed using a Qubit dsDNA HS assay kit (Life Technologies, # Q32854) and Qubit spectrometer (Invitrogen #Q33226) of 12 wells of cDNA / plate, with cDNA from each of the four biological samples represented on the plate, as per the manufacturer’s guidelines. Each plate was diluted based on a dilution factor for a median concentration of input cDNA ~ 200ng/μL.

Nextera XT tagmentation and adaptor ligation was performed as described in the SMART-Seq HT manual, using the Nextera XT DNA Library Preparation Kit (Illumina, #FC-131-1096) and the Nextera Index Kit v2 Set A (Illumina, #FC-131-2001). Aliquots of the tagmented and adaptor ligation PCR product were pooled (Each 96 well plate was pooled into a single 1.5mL microcentrifuge tube), and the pool was cleaned with AMPure XP beads as described in the SMART-Seq HT manual, using the 1.5mL centrifuge tube magnetic stand (Takara, #621964). The final library concentrations and qualities were determined by both SYBR PCR (Kapa Biosystems, #KK4824) with a BioRad CFX qPCR machine and Bioanalyser (High Sensitivity DNA assay, Agilent). Libraries were sequenced on an Illumina NextSeq 500 with 75bp paired end reads and a mid-output run (96 cells/run), yielding approximately one million aligned reads / cell.

### Gene expression analysis and quality control of single cell SMART-Seq data

Reads from demultiplexed fastq files were trimmed with trimgalore (v 0.6.6) and aligned to the human genome (Ensembl build GRCh38.87) using HISAT2 (v 2.1.0, (Kim et al., 2015)) with the options --sp 1000,1000. Aligned reads were quantified using *featureCounts*, as implemented in *Rsubread* (Subread v 2.0.0, (Liao et al., 2019)), and R 3.6.1. Downstream analyses were performed using Bioconductor (Huber et al., 2015) packages: *scater* (McCarthy et al., 2017), *scran* (Lun et al., 2016), and the *SingleCellExperiment* (Lun and Risso, 2019) data infrastructure, as outlined here (Amezquita et al., 2020).

Gene expression data was obtained for 952 cells. The following quality control steps were applied, within each biological condition (22-36 yo d0; 22-36 yo d42; 67-86 yo d0; 67-86 yo d42), excluding cells meeting one or more of the following criteria: i. the number of detected transcripts >3x median absolute deviations (MAD) from the median ii. the percentage of reads that map to mitochondrial transcripts >3x MAD from the median iii. the percentage of the library that is occupied by the top 50 transcripts (a measure of low complexity libraries) > 3x from the median (Supplementary Figures 2A-D). 117 samples were excluded for low library size (Supplementary Figures 2A). 130 samples were excluded for low numbers of detected transcripts (Supplementary Figures 2B). 36 samples were excluded for high proportions of mitochondrial transcripts (Supplementary Figures 2C). 119 samples were excluded for low complexity libraries (Supplementary Figures 2D). Cells could fail on several QC parameters. The total number of cells discarded was 163. Initially, Highly variable genes (HVGs) were identified as the top 25% of variance, after normalization for library size using deconvolution. The first 50 components from principal component analysis of these HVGs were used for UMAP embedding (McInnes et al., 2020). Clustering was performed with Louvain clustering in *igraph* (Csardi and Nepusz). After excluding cells on biological grounds (see Supplementary Figures 1F-K), a new set of HVGs were defined, using the top 10% of variance and the first 40 principal components for UMAP. Cell assignment was performed with *SingleR* (Aran et al., 2019), and the Monaco *et al*. dataset (Monaco et al., 2019). For clarity, particularly around double-negative B cells, we have re-labelled the Monaco *et al*. B cell populations as follows [sorting strategy, label herein]: IgD^+^CD27^-^, naïve; IgD^+^CD27^+^, non-switched Bmem; IgD^-^CD27^+^, switched Bmem; IgD^-^CD27^-^, IgD^-^CD27^-^ Bmem; IgD^-^CD27^++^CD38^+^, plasmablasts. Trajectory analysis was performed using the *destiny* package (Angerer et al., 2015). Differential abundance analysis was performed using *edgeR* (Robinson et al., 2010).

### B cell receptor assembly from single cell SMART Seq data

We used VDJPuzzle (Rizzetto et al., 2018) to retrieve and annotate heavy and light chains from each individual B cell. This gives, per cell, annotated heavy chain and light chain alignments, and a report of detected mutations. Amino acid positions follow the Kabat numbering scheme. Where VDJPuzzle reported more than one chain, we report the chain with the highest expression level (for example, in ~50% of naïve cells, identical V, D and J segments were assembled with an IgM or IgD heavy chain). For replacement:silent ratios of somatic hypermutation, these are pseudolog transformed prior to plotting. The pseudolog distribution becomes linear at around 1 and allows both the benefits of a log compression of large numbers and, unlike a log transformation, the retention of cells where R/S+0.01=0, due to R=0. These analyses on the VDJPuzzle results were performed in R 3.6.1.

### Gene expression analysis and B cell receptor analysis from 10x data

The published Turner *et al*. dataset (Turner et al., 2020) was downloaded as fastq from ENA (https://www.ebi.ac.uk/ena/browser/home), using aspera links retrieved from the SRA Explorer (http://sra-explorer.info/). Fastq files missing from ENA were re-created using fastq-dump from the SRAtoolkit (v2.10.8, http://ncbi.github.io/sra-tools/). Gene expression data was aligned and counted using cellranger count (v5.0.0, 10x Genomics) against the human genome (10x Genomics build GRCh38-1.2.0). Downstream analysis was performed with the *Seurat* R package (Stuart et al., 2019). Quality control retained cells with: >200 features/cell & <7,000 features/cell & < 12.5% mitochondrial reads. Each sample was then normalized and the 2000 most variable features selected, after variance stabilized normalization. For either FNA or IgD-PBMC samples, all time points were combined using *ImmuneAnchors* to minimize batch effects between replicates (first 30 dimensions used). To identify B cells with FNA or IgD-PBMC samples, we used 20 principal components for clustering, and *FindClusters* with a resolution of 0.05. B cells were selected on the basis of the expression of *CD79A, MS4A1* and *CD19* (Turner et al., 2020). Once identified, B cells were re-scaled and re-clustered with 20 principal components (*FindClusters* resolution = 0.05). GC B cells were identified as the FNA B cell cluster expressing *BCL6, RGS13, MEF2B, STMN1, ELL3, SERPINA9* (Turner et al., 2020). Resting memory B cells were defined as B cells expressing *TNFRSF13B*, *CD27* and *CD24* (Turner et al., 2020).

VDJ repertoire data were pre-processed using cellranger vdj (v5.0.0, 10x Genomics), with the VDJ annotations from 10x Genomics (refdata-cellranger-vdj-GRCh38-alts-ensembl-5.0.0). Mutation analysis was performed with the Immcantation suite (Gupta et al., 2015), and replacement:silent ratios calculated and plotted as described above for the SMART-Seq dataset. Clonotype tracking was performed using the *immunarch* R package (Vadim Nazarov et al., 2020). Clonotypes were defined having identical *IGHV, IGHJ, IGLV* and *IGLV* alleles, and IgH and IgL CDR3 lengths.

### Statistical analyses

All statistical analyses were performed in R (v 3.6.1, www.r-project.org) via RStudio (v1.2.5001; www.rstudio.com). The statistical test used is indicated in the relevant figure legend. Figures were generated using Rmarkdown (Allaire et al., 2021; Xie et al., 2018, 2020) with pdf output.

## Code and data availability

Corresponding R code for the SMART-Seq gene expression and BCR analyses, 10x/Seurat analyses, and phenotypic data is available at the github repository: https://github.com/lintermanlab/B_flu_scRNAseq. This repository includes Rmarkdown files to reproduce each figure. Raw single cell sequencing data are deposited in GEO (https://www.ncbi.nlm.nih.gov/geo/) under the accession GSE167823. The Turner *et al*. dataset (Turner et al., 2020) is available from GEO with the accession GSE148633. Other primary data, for example raw FCS files, are available from the authors upon reasonable request.

## Supplemental material

Supplementary Figure 1 shows the full gating for single cell sorting and calculations of the abundance of haemagglutinin specific B cells in PBMC. Supplementary Figure 2 shows scRNAseq quality control metrics and the identification of a plasmablast population. Supplementary Figure 3 shows the transcriptional heterogeneity of haemagglutinin specific B cells. Supplementary Figure 4 relates to Figure 5, showing additional steps in the definition of B cells in FNA and IgD-enriched PBMC, and GC B cells. It also shows the SHM profiles of circulating *FCRL5*^+^ memory B cells.

## Author contributions

EJC designed the study, performed experiments, analyzed data, performed bioinformatics analyses, wrote the manuscript, obtained funding and provided clinical oversight.

AKW provided reagents.

DLH performed experiments, analyzed data and wrote the manuscript.

MAL designed the study, performed experiments, wrote the manuscript, obtained funding and oversaw the project.

The authors declare no competing interests.

## Acknowledgements

The NIHR Cambridge Biomedical Research Center (BRC) is a partnership between Cambridge University Hospitals NHS Foundation Trust and the University of Cambridge, funded by the National Institute for Health Research (NIHR). We are indebted to the NIHR Cambridge BRC volunteers for their participation and we thank the NIHR Cambridge BRC staff for their contribution in co-ordinating the vaccinations and venepuncture. We thank the staff of the Babraham Institute Flow Cytometry Facility, the Sequencing Facility, and Silvia Innocentin for their technical assistance. We are grateful to the Babraham Bioinformatics Group for their insightful discussions. We thank Dr Geoff Butcher for his comments on this project throughout its conception and delivery.

This study was supported by H2020 European Research Council funding awarded to MAL (637801-TWILIGHT), Evelyn Trust funding (19/20) awarded to EJC and MAL and the Biotechnology and Biological Sciences Research Council (BBS/E/B/000C0427, BBS/E/B/000C0428, and the Campus Capability Core Grant to the Babraham Institute). DLH is supported by a National Health and Medical Research Council Australia Early-Career Fellowship (APP1139911). MAL is an EMBO Young Investigator and Lister Institute Prize Fellow.

This paper presents independent research supported by the NIHR Cambridge BRC. The views expressed are those of the authors and not necessarily those of the NIHR or the Department of Health and Social Care.

---

**Supplementary Figure 1:**
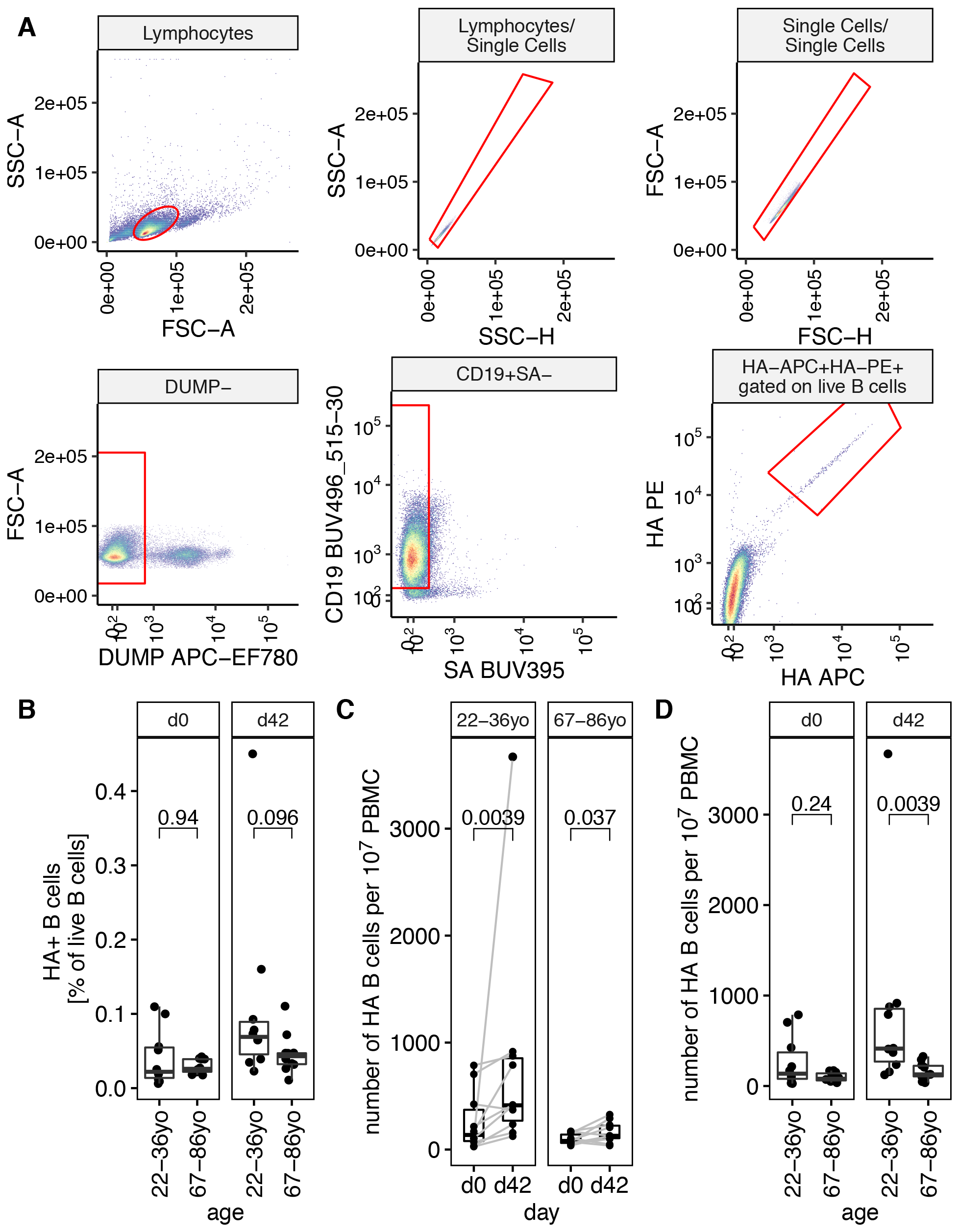
Single cell sorting strategy for hemagglutinin specific B cells and abundance of haemagglutinin specific B cells. (A) Example flow sorting gating strategy on B cells. B cells were negatively separated from PBMC using magnetic sorting prior to flow sorting. (B) The proportion of haemagglutinin binding B cells (as % of live B cells, that did not bind free-streptavidin), is not significantly different between age groups either before, or 6 weeks after TIV immunization. (C-D) The number of haemagglutinin binding B cells per 10^7^ PBMC analysed by age and days post vaccination. In (B) and (D), the *P* values shown are from an unpaired two-tailed Mann-Whitney test. In (C), samples from the same individual are indicated with a grey line. The *P* values shown are from a paired two-tailed Mann-Whitney test.

**Supplementary Figure 2:**
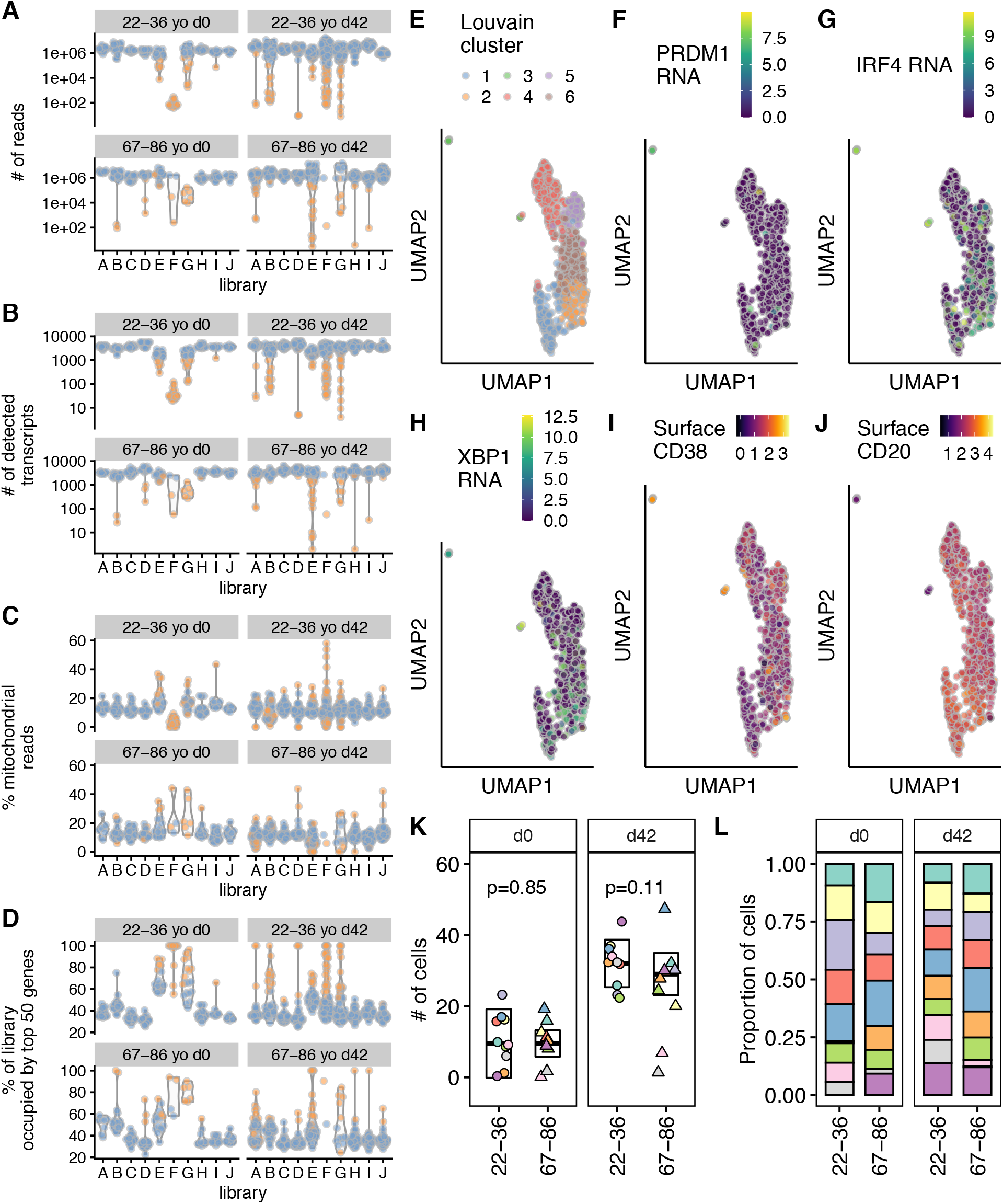
Single cell RNA sequencing quality control. (A) The number of reads per cell (n=952) plotted on a log10 scale for each library, with the 4 experimental conditions (22-36 year old at day 0; 22-36 year old at day 42; 67-86 year old at day 0; 67-86 year old at day 42) shown separately. (B) The number of detected transcripts per cell plotted as in (A) plotted on a log10 scale for each library, with the 4 experimental conditions shown separately. (C) The percentage of detected mitochondrial transcripts per cell plotted as in (A). (D) The percentage of reads per cell that are attributable to the 50 most highly expressed features. Cells with high values represent low complexity libraries. (E) UMAP plot of transcriptomes of single HA-binding B cells, after QC, n=789 cells. The first 50 principal components were calculated with features selected by the top 25% of variance, after normalization for library size using deconvolution. For (F)-(J) UMAP embedding as in (E), with the following: (F) Expression level of *PRDM1* (G) Expression level of *IRF4* (H) Expression level of *XBP1* (I) Logicle fluorescence intensity of surface CD38 protein (J) Logicle fluorescence intensity of surface CD20 protein (K) The number of cells, that pass transcriptomic QC, and are not plasma cells (E-J) for 22-36yo and 67-86yo individuals at day 0 and day 42; n=771 cells. The median and median absolute deviations, and *P* values from 2 tailed Mann-Whitney tests are shown. Each individual is identifiable by their plotted shape and its color. There is 1 22-36yo individual (purple circle), who only provides cells only at day 42 and one 67-86yo individual (pink triangle) who only provides cells at day 42. (L) The percentage of each condition’s cells, after transcriptomic QC, that each individual contributes is plotted.

**Supplementary Figure 3:**
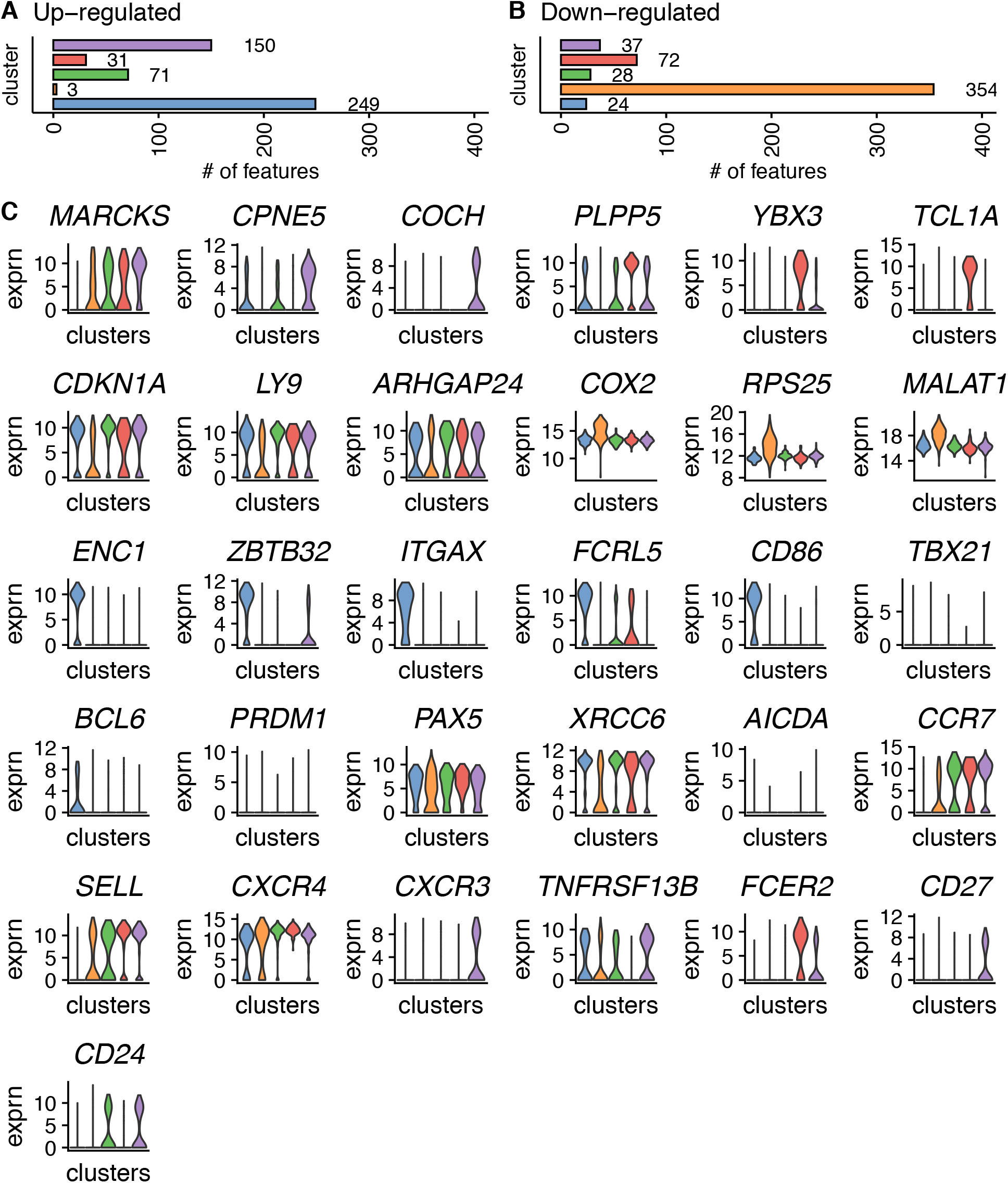
Transcriptional heterogeneity of haemagglutinin specific B cells. (A) Up-regulated differentially expressed transcripts between a cluster and *any* other cluster, where log2 fold change > 2 and Benjamini-Hochberg FDR<0.01, using pairwise t-tests. For each cluster the number of markers is indicated. (B) As in (A), for down-regulated markers. (C) Violin plots of gene expression of the indicated gene for each of the 5 UMAP clusters. Expression (log-transformed normalized expression) values are plotted. Colors of each UMAP cluster are the same as in (A), (B) and the main figures.

**Supplementary Figure 4:**
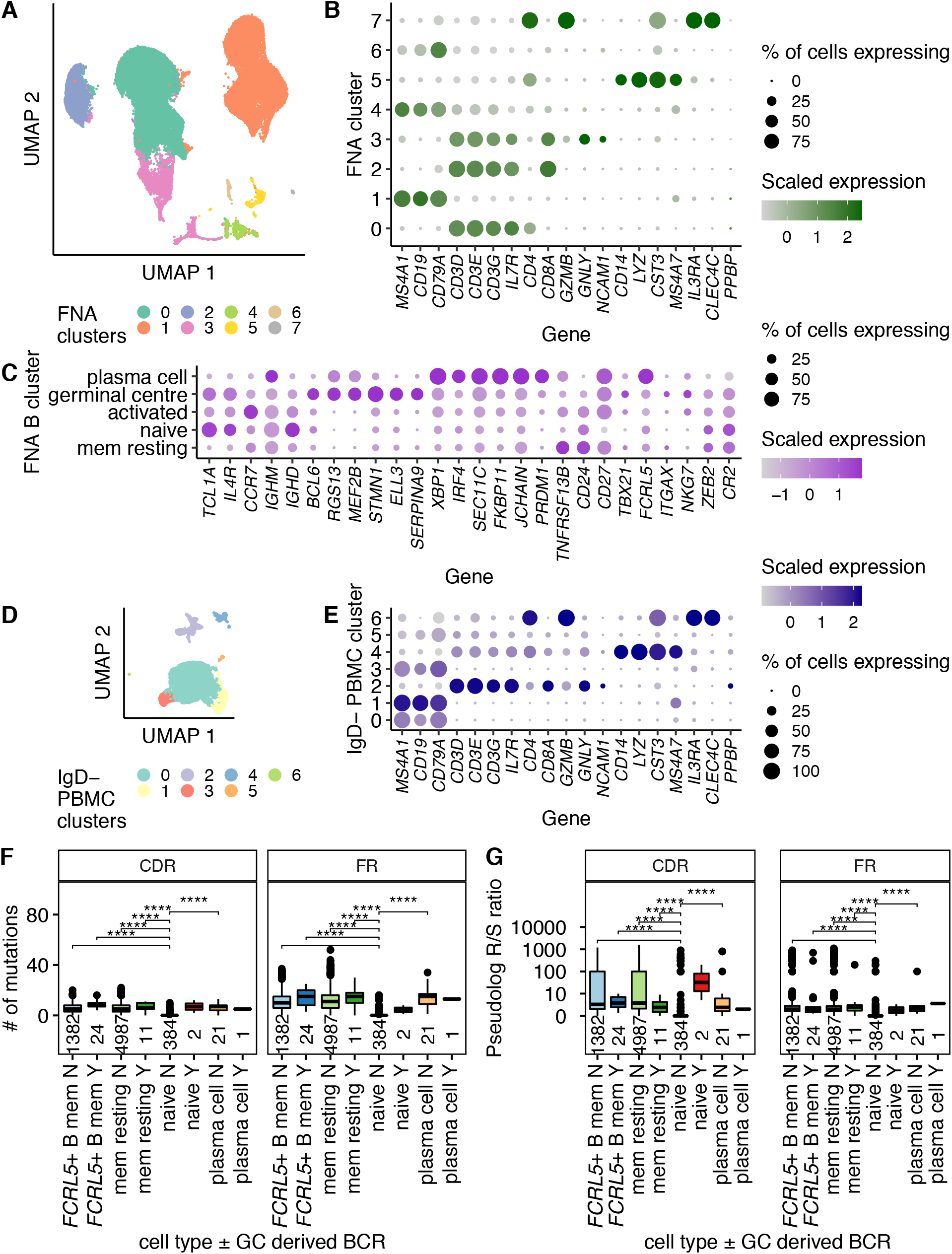
Defining B cells within fine needle aspirations and B memory enriched PBMC samples, and somatic hypermutation rates of B cell subsets. (A) UMAP of all lymph node fine needle aspiration cells (n=70,795), with Louvain clusters shown. (B) Identification of B cell clusters in (A), based on the expression of *CD19, MS4A1* and *CD79A*. Other key lineage markers, as used in Turner *et al*. (Turner et al., 2020), are shown: T cells *CD3D, CD3E, CD3G, IL7R, CD4, CD8A*; NK cells *GZMB, GNLY, NCAM1*; monocytes *CD14, LYZ, CST, MS4A7*; plasmacytoid dendritic cells *IL3RA, CLEC4C* and platelets *PPBP*. (C) Identification of the fine needle aspirate B cell sub-clusters in Figure 5A, based on the expression of markers as used in Turner *et al*. (Turner et al., 2020): naïve B cells *TCL1A, IL4R, CCR7, IGHM, IGHD*; germinal center B cells *BCL6, RGS13 MEF2B, STMN1*, *ELL3*, *SERPINA9*; plasma cells *XBP1*, *IRF4*, *SEC11C*, *FKBP11*, *JCHAIN*, *PRDM1*; resting memory B cells (mem resting) *TNFRSF13B, CD27* and *CD24*; activated B cells *TBX21, FCRL5, ITGAX, NKG7, ZEB2*, and the lack of *CR2*. (D) UMAP of all Bmem enriched (IgD-) PBMC (n=24,156 cells), with Louvain clusters shown. (E) Annotation of B cell subsets in (D), based on the expression of marker genes as in (B). (F) The number of mutations in the CDR or FR regions of B cell receptors for each B cell subset, from B cells that share a BCR (Y) with the day 12 germinal center and those that do not (N). The number of cells in each category is shown just above the horizontal axis. (G) The replacement:silent ratios for the CDR or FR regions of B cell receptors for each B cell subset, from B cells that share a BCR (Y) with the day 12 germinal center and those that do not (N). B cell subsets are defined in Figures 5C & D. The number of cells in each category is shown just above the horizontal axis. As in Figure 4, the replacement:silent ratio is calculated as # of replacement mutations / (# of silent mutations + 0.01), to avoid discarding cells with zero silent mutations. The resulting R/S ratio is plotted as a pseudolog. P values shown are from two-tailed Mann-Whitney tests, comparing the indicated cluster against naïve cells.

